# PDF neuropeptide signals independently of Bruchpilot-labelled active zones in daily remodelled terminals of *Drosophila* clock neurons

**DOI:** 10.1101/2023.06.20.545701

**Authors:** Benedikt Hofbauer, Meet Zandawala, Nils Reinhard, Dirk Rieger, Christian Werner, Jan-Felix Evers, Christian Wegener

## Abstract

The small ventrolateral neurons (sLNvs) are key components of the central clock in the *Drosophila* brain. They signal via the neuropeptide Pigment-dispersing factor (PDF) to align the molecular clockwork of different central clock neurons and to modulate downstream circuits. The dorsal terminals of the sLNvs undergo daily morphological changes that have been shown to affect presynaptic sites organised by the active zone protein Bruchpilot (BRP), a homolog of mammalian ELKS proteins. Although the circadian plasticity of the sLNv terminals is well established, whether and how it is related to the rhythmic release of PDF remains ill-defined.

Here, we combined expansion microscopy with labelling of active zones by endogenously tagged BRP to examine the spatial correlation between PDF-containing dense-core vesicles and BRP-labelled active zones. We found that the number of BRP-labelled punctae in the sLNv terminals remained stable while their density changed during circadian plasticity. The relative distance between BRP- and PDF-labelled punctae was increased in the morning, around the reported time of PDF release. Spontaneous dense-core vesicle release profiles of the sLNvs in a publicly available ssTEM dataset (FAFB) consistently lacked spatial correlation to BRP-organised active zones. RNAi-mediated downregulation of *brp* and other active zone proteins expressed by the sLNvs did not affect PDF-dependent locomotor rhythmicity. In contrast, down-regulation of genes of the canonical vesicle release machinery, the dense-core vesicle-related protein CADPS, as well as PDF impaired locomotor rhythmicity.

Taken together, our study suggests that PDF release from the sLNvs is independent of BRP-organised active zones which seem not to be circadianly destroyed and re-established.

## Introduction

Pigment-dispersing factor (PDF) is arguably the best characterised insect “circadian” neuropeptide. PDF serves a dual signalling function by aligning the molecular clockwork of the different central clock neurons (Liang *et al*., 2016, 2017) and serving as a major clock output that modulates various downstream circuits (Shafer & Yao, 2014; King & Sehgal, 2018). Within the *Drosophila* clock network, PDF is expressed in a small set of small and large ventrolateral neurons (sLNvs and lLNvs, respectively) that are referred to as circadian pacemaker neurons (Helfrich-Förster, 1997; Shafer & Yao, 2014). Both LNv types as well as PDF are required for normal locomotor rhythmicity (Renn *et al*., 1999; Shafer & Yao, 2014). In addition to PDF, the sLNvs also express short neuropeptide F (sNPF (Johard *et al*., 2009)) and the classic neurotransmitter glycine (Frenkel *et al*., 2017).

The sLNvs have dendritic processes in the accessory medulla, and rhythmically release PDF from axon terminals in the superior protocerebrum during the beginning of the light phase (Park *et al*., 2000; Klose *et al*., 2021). This release is dependent on membrane excitability (Nitabach *et al*., 2006). Concomitantly, the sLNvs show significantly higher electrical activity in the morning than in the evening (Cao *et al*., 2013), and intracellular free Ca^2+^ levels that peak at the end of the night and decline over the day (Liang *et al*., 2017).

Intriguingly, the dorsal terminals of the sLNvs undergo significant time-dependent morphological changes, with maximum branching and complexity in the early morning (Zeitgeber time ZT2) compared to early (ZT14) or late night (Fernández *et al*., 2008; Gorostiza *et al*., 2014). This time-dependent plasticity depends on temporal changes in membrane activity (Depetris-Chauvin *et al*., 2011; Sivachenko *et al*., 2013; Gorostiza *et al*., 2014) and is driven by the circadian clock as it persists in constant darkness (DD) but is lost in flies without a functional clock (Fernández *et al*., 2008). While the underlying mechanisms are not entirely clear, Fasciclin 2 (Fas2)-dependent axonal de- and re-fasciculation and Rho1-dependent axonal retraction are known to be involved (Sivachenko *et al*., 2013; Petsakou *et al*., 2015). PDF signalling is additionally required for the diel morphological plasticity, as manipulating PDF signalling or matrix metalloproteinases that degrade PDF affects the structural plasticity (Depetris-Chauvin *et al*., 2014).

The dorsal terminals of the sLNvs contain both presynaptic and (sparser) postsynaptic sites (Yasuyama & Meinertzhagen, 2010) and can be labelled by a dendritic marker (Petsakou *et al*., 2015), suggesting that they represent a mixed in- and output compartment. At least the presynaptic sites seem to undergo a daily change in numbers, in correlation with the morphological remodelling. In *Drosophila,* the presynaptic active zone is organised by Bruchpilot (BRP), a fly homolog of the vertebrate ELKS/CAST active zone proteins that clusters presynaptic Ca^2+^ channels (Kittel *et al*., 2006; Wagh *et al*., 2006; Fouquet *et al*., 2009; Ghelani & Sigrist, 2018). Ectopic *pdf*-Gal4-mediated expression of a tagged BRP version (BRP^RFP^) in the sLNvs revealed a significantly higher number of BRP^RFP^-positive punctae at circadian time (CT) 2 compared to CT14 and CT22 (Gorostiza *et al*., 2014). Moreover, the number and total area of punctae labelled by a Synaptotagmin-GFP reporter (SYT^GFP^, labels both peptidergic dense-core as well as synaptic vesicles) in *pdf>Syt^gfp^* flies was increased at CT2 compared to CT14 (Gorostiza *et al*., 2014). Activity-dependent GFP Reconstitution Across Synaptic Partners (GRASP) further points towards a time-dependent switch of postsynaptic partners of the sLNvs (Depetris-Chauvin *et al*., 2014). Collectively, these findings suggest that the circadian plasticity of the sLNv terminals is coupled to a change in synapse number, synaptic contacts, and active zones of synaptic vesicle release. This led to a model in which the diel morphological and synaptic changes contribute to rhythmic peptidergic and transmitter output of the sLNvs which in turn drives downstream circuits regulating rhythmic behaviour and physiology. In support of this idea, overexpression of *Rho1,* which locks the terminal arborisations in a nocturnal state, leads to arrhythmic locomotor activity in constant darkness (DD (Petsakou *et al*., 2015)), a canonical phenotype of impaired PDF signalling.

A recent study, however, genetically prevented the formation of the dorsal sLNv terminals and found that sLNv/PDF-mediated circadian output functions are unaffected (Fernandez *et al*., 2020). The PDF-dependent (Renn *et al*., 1999) bimodal diel pattern of locomotor activity as well as circadian rhythmicity and resetting of other clock cells in DD was normal in flies lacking the dorsal sLNv terminals (Fernandez *et al*., 2020). These findings suggest that rhythmic PDF release from the sLNvs does neither require the dorsal terminals nor their diel plasticity. Moreover, at least in isolated brains, PDF can also be released from the sLNv somata, albeit with a phase different from terminal PDF release (Klose *et al*., 2021). These findings raise the question to which extent PDF release requires BRP-organised presynaptic active zones in the dorsal terminals.

Here, we anatomically examined the spatial correlation between PDF-containing dense-core vesicles and release sites and BRP-organised active zones in the sLNvs at the ultrastructural level using super-resolution fluorescence and electron microscopy. We further performed a targeted RNA interference (RNAi)-based screen to test the functional relevance of sLNv-expressed structural active zone proteins and proteins of the vesicle fusion machinery for PDF signalling, using locomotor activity as a behavioural read-out. Our results suggest that PDF release from the sLNv is largely independent of the presence and localisation of BRP but requires SNARE and vesicle fusion proteins. The independence of PDF release from BRP-organised active zones helps to explain the previously reported PDF release from sLNv somata and abrogated dorsal terminals(Fernandez *et al*., 2020; Klose *et al*., 2021). Our results further suggest that BRP-labelled puncta remain stable during the day and become compacted during circadian terminal retraction at the beginning of the dark phase.

## Material and Methods

### Fly husbandry

The following genotypes of *Drosophila melanogaster* were used for synaptic labelling: UAS-brp::GFP-TR725 (BL36292 (Fouquet *et al*., 2009)), dFLEX w; *Pdf*-Gal4/BrpFOn Ypet, UAS-FLP; +/MKRS (Gärtig *et al*., 2019). The lines for the targeted RNAi screen are listed in Suppl. Table 1 and originated from the shRNA, KK, and GD library of the Vienna Drosophila Resource Center (VDRC) or were a kind gift of Stephan Sigrist (Freie Universität Berlin, Germany), Paul Taghert (U Washington, St. Louis, USA) or Michael Bender (U Georgia, Athens, GA, USA).

Flies were maintained under LD 12:12 and 60 ± 5% humidity at 18± 0.2 °C or 25± 0.2 °C on standard medium containing 8.0% malt extract, 8.0% corn flour, 2.2% sugar beet molasses, 1.8% yeast, 1.0% soy flour, 0.8% agar and 0.3% hydroxybenzoic acid.

### Immunocytochemistry (ICC)

Brains were dissected under a stereoscopic microscope and pooled in reaction tubes with 0.1 M phosphate-buffered saline (PBS, pH 7.4) on ice, followed by fixation in 2% paraformaldehyde and 4% sucrose in PBS for 1 hour at RT. After at least 5x 5 minutes washing steps with PBS, the tissues were blocked with 5% normal goat serum (NGS) in PBS containing 0.3% Triton-X100 (PBT) for 2 hours at RT. Then, the brains were incubated in primary antibody (mouse monoclonal anti-PDF C7, 1:500-1:2000, DHSB, donated by Justin Blau; rabbit polyclonal anti-GFP 1:1000-2000, ChromoTek) solution in PBT+5% NGS at least overnight at 4°C, followed by 6x 5 minutes washes at RT with PBT. This was followed by incubation in secondary antibody (goat anti-rabbit IgG coupled to Alexa Fluor 488 or goat anti-mouse IgG coupled to Alexa Fluor 568, 1:500, Invitrogen) solution in PBT+5% NGS for 3 hours at RT or over-night at 4°C, and washes for 4x 10 minutes in PBT and 2 x 10 minutes in PBS at RT. The brains were then moved into a PBS droplet on a high precision coverslip (Marienfeld, 22×22mm) with two hole-reinforcement rings (LEITZ) as spacer. The PBS was then removed, and tissues were mounted in Vectashield (Vector Laboratories) and sealed with a second coverslip on top. Samples were stored at 4°C for a few days before scanning.

### Whole Mount Microscopy and Image Analysis

Preparations were scanned on a Leica TCS SP8 confocal laser-scanning microscope (Leica Microsystems, Wetzlar, Germany) equipped with hybrid detectors, photon multiplier tubes, and a white light laser for excitation, together with a 20-fold and 63-fold glycerol immersion objective (HC PL APO, Leica Microsystems, Wetzlar Germany). The acquisition parameters were set to saturation of the signal and were maintained constant throughout the scanning process. Images were processed and analysed in Fiji (Schindelin *et al*., 2012). To produce the figures maximum projections were compiled and brightness and contrast were linearly adjusted. The PDF signal was used to create a binary mask for each image-stack, by manually enhancing the saturation of the PDF channel before setting an arbitrary threshold to create a binary mask-stack of the neuron. The mask was used to capture BRP signal that was co-located with PDF. Hand-defined masks were further utilized to define compartments of the neurons: dendrites, kink, and axon. A custom-written simple Fiji macro was employed to quantify and measure the BRP particles that were included in the binary PDF mask. Mann-Whitney-U-Test in Python (SciPy-library) was used for statistical analysis.

### Tissue expansion

For expansion microscopy (ExM, (Chozinski *et al*., 2016)), ICC was performed as described above, with a few additional steps as described earlier (Gärtig *et al*., 2019). After ICC, samples were transferred from the reaction tubes into a droplet of PBS on poly-Lysine coated glass slides (ThermoFisher) stabilised by a bean-shaped ring drawn with a PAP-Pen (Merck). The slides were then transferred to a moist chamber. Under the fume hood, PBS was replaced by 1mM methacrylic-acid N-hydroxysuccinimide-ester (MA-NHS) in PBS for 1 hour at RT and then washed with PBS (3x 5 minutes). Afterwards, tissues were incubated first in 30%, then 60% monomer solution (MS, 38% sodium acrylate, 40% acrylamide, 2% N,N’-methylenebisacrylamide, 29.2% sodium chloride, filled up by PBS to 10 ml) for 10 minutes, and then 100% MS for 30 minutes at 4°C. MS was then replaced by 80µl of fresh 100% MS solution and a gelling chamber (coverslip with glass spacers) was placed on top. The MS solution was then removed and gelling solution (0.01% 4-hydroxy-TEMPO, 0.2% Tetramethylethylenediamine (TEMED) and 0.2% ammonium persulfate in MS) was added in parallel. Gelling occurred for 2-2.5 hours at 37°C.

After gelling, the chamber was carefully removed, and excess gel was cut away. To facilitate later orientation, the samples were cut into a trapezoid shape, and transferred into a 6-well plate with 800µl digestion buffer (50 mM Tris pH 8.2, containing 1 mM EDTA, 0.5% Triton X-100, 0.8 M guanidine-HCl, 8 units/ml Proteinase K, ddH_2_O added to 5 ml) and digested for 2 h at 37°C. After digestion, the gels were washed with an excess volume of distilled water (5 x 10 minutes).

We also applied a post-expansion staining protocol to increase the fluorescence strength of the samples. The protocol is similar to the standard expansion protocol above but uses a different linker and adds another staining step after the digestion and the initial expansion step. PBS was replaced by 1 mM acryloyl X-SE ester (AcX), and tissues were incubated overnight at RT and then washed with PBS (3x 15 minutes). After digestion, the distilled water was replaced by PBS, and the gels were incubated in primary antibody solution (5% NGS in PBS) at the same concentrations as for ICC described above at 4°C overnight. Gels were then washed with PBS (6x 10 minutes) and incubated in secondary antibody solution as for ICC described above for at least 3 hours. Next, gels were washed two times with PBS, and then incubated in 0.1% Hoechst 33342 for 30 min. Finally, gels were washed again with PBS (2 x 10 minutes) and then expanded in distilled water (5x 10 minutes).

### Expansion microscopy (ExM)

After expansion resp. post-expansion staining, the gels were transferred onto a glass slide with a silicon spacer chamber on top. The chamber around the gel was filled with 2% low melt agarose and sealed by a coverslip taped on top. This allowed imaging by the Leica TCS SP8 confocal laser-scanning microscope described above.

### Image Analysis

Regions of interests (ROIs) were defined by PDF staining by applying a soft focus and creating a binary mask. The binary mask was then used as a template to extract BRP puncta that were located close to PDF staining. The number and size of BRP and PDF-labelled puncta were then quantified using the “analyse particles” function of ImageJ. To analyse the distance between BRP and PDF puncta, the channels of each staining were transformed into binary masks. A 3D distance map was created based on the BRP mask and combined with the PDF binary mask to determine the relative distance of PDF particles from BRP staining as gray values. The resulting distance map was processed with the PDF-mask containing the values of Euclidean distance between BRP puncta for each PDF particle as defined by the PDF mask and analysed with the “analyse particles” function to collect the mean value for each PDF punctum. Statistical analysis of the expansion images was performed with the Mann-Whitney-U-Test in Python (scipy-library).

To quantify the density of BRP and PDF puncta in the terminals, the number of respective puncta was divided by the area of the binary mask for each section of the image stack. The median of the results was calculated for each image stack and plotted.

#### CATMAID

To utilize the huge dataset of the full adult female fly brain (TEMCA2) (Zheng *et al*., 2018), the 12 TB dataset was set up locally on an instance to enable local tracing in CATMAID (Saalfeld *et al*., 2009). Based on the characteristic structure of the PDF-positive sLNvs and experience from immunostained ultrastructural samples, a set of seven neurons (3 resp. 4 on each hemisphere) was identified in the TEMCA-dataset that showed close morphological resemblance to the sLNv structure and contained DCVs. These putative s-LNvs were completely traced by hand and the ultra-structure was manually analysed for DCV fusion events with the cell membrane in the dorsal area. The skeletons were retrieved from the local CATMAID server using the CATMAID-to-Blender Plugin (v7.0.2, (Schlegel *et al*., 2016)) for Blender (v3.5.0, blender.org). Omega fusion sites of DCV within the putative s-LNvs were annotated by hand in CATMAID and the coordinates plotted using the rgl libraries (v1.0.1 (Murdoch & Adler, 2023)) in RStudio (v2022.12.0, Posit Software, PBC) for R (v4.2.2, R-project.org). The s-LNv neuron meshes from the hemibrain dataset (v1.2.1, (Scheffer *et al*., 2020)), and the automatic annotated presynaptic sites (Buhmann *et al*., 2021) were obtained from the neuprint server (neuprint.janelia.org) using the natverse libraries (v0.2.4, (Bates *et al*., 2020)) in RStudio. The neuron meshes and coordinates of the presynaptic sites were transformed to the FAFB coordinate space (FAFB14) for better comparison, using xform_brain (nat.templatebrains v1. 0). The final visualisation was performed in Blender.

### Locomotor activity monitoring

Locomotor activity was monitored using commercial *Drosophila* Activity Monitors (DAM-2, Trikinetics, Waltham, USA). 3-5 d old male flies were individually placed in small glass tubes (5 mm diameter) which contained food on one side (2% agar containing 4% sucrose) closed with a rubber plug. The other side was closed with a foam plug to allow for air exchange. Monitors were placed into light boxes manufactured by our workshop. Activity of the flies was measured by counting the interruptions of the infrared beam in the middle of the tubes. White light was supplied at 1 (LL) or 100 (LD) lux. Activity was monitored at a constant temperature of 20°C in LD12:12 or LD16:8 conditions for 6 days, followed by constant darkness (DD) or constant light (LL) for at least a week (DD1–10 or LL1-10).

Double-plotted actograms were generated by a script in R provided by Enrico Bertolini (Bertolini *et al*., 2019). Activity was summed into 15-minute bins. Rhythmicity parameters were analysed by autocorrelation with a MATLAB script developed by Joel Levine and colleagues (Levine *et al*., 2002). Flies were scored as arrhythmic if their rhythmic strength after autocorrelation was below 15. The percentage of rhythmic flies was calculated as the number of rhythmic flies divided by the total number of flies that survived for at least 10 days under constant conditions. Data distribution was tested for normality with the Shapiro-Wilk-Test and significance was calculated via the Mann-Whitney-U-Test in Python.

### Expression analysis of synaptic and vesicle release-associated proteins

Gene transcripts coding for synaptic or vesicle-release associated proteins in clock neurons (sLNvs and lLNvs) were mined in previously published single-cell RNA sequencing (scRNA-seq) datasets (Ma *et al*., 2021; Li *et al*., 2022). Pre-processing, dimensionality reduction, and clustering of clock cell scRNA-seq data was performed using the original code (Ma *et al*., 2021). All analyses were performed with the Seurat package (v4.1.1 (Hao *et al*., 2021)) in R-Studio (v2022.02.0). Heatmaps were generated using the pheatmap package (v1.0.12).

## Results

### Intragenic BRP markers reliably label active zones when expressed at endogenous levels

Previous work used *Pdf*-GeneSwitch>UAS-mediated ectopic expression of BRP^RFP^ to label active zones and to quantify their temporal changes (Gorostiza *et al*., 2014). We adopted a similar strategy by using *Pdf*-Gal4 to drive UAS-*brp::GFP* (Fouquet *et al*., 2009) in the sLNvs. We successfully labelled punctate structures along the whole proximal axon, the lateral flexure in the superior protocerebrum (“kink”) and the more median dorsal terminals in the superior protocerebrum (Fig. 1A-B) confirming earlier findings (Gorostiza *et al*., 2014). Surprisingly, the size of the BRP^GFP^-labelled punctae was unusually large for active zones (around 1 µm at confocal resolution (Fig. 1B)). Furthermore, BRP^GPF^-labelling was not localised to the plasma membrane as expected but was localised more luminal than PDF co-immunolabelling indicating assemblies of peptidergic dense-core vesicles (Fig 1B). In addition, the cell bodies were strongly labelled by BRP^GFP^. The observed unusual size and location of BRP^GFP^ labelling in the sLNvs suggests that, at least in our hands, *Pdf*-Gal4-mediated overexpression of BRP leads to protein aggregation of BRP^GFP^ and is not a good reporter for the distribution of native BRP.

**Figure 1:**
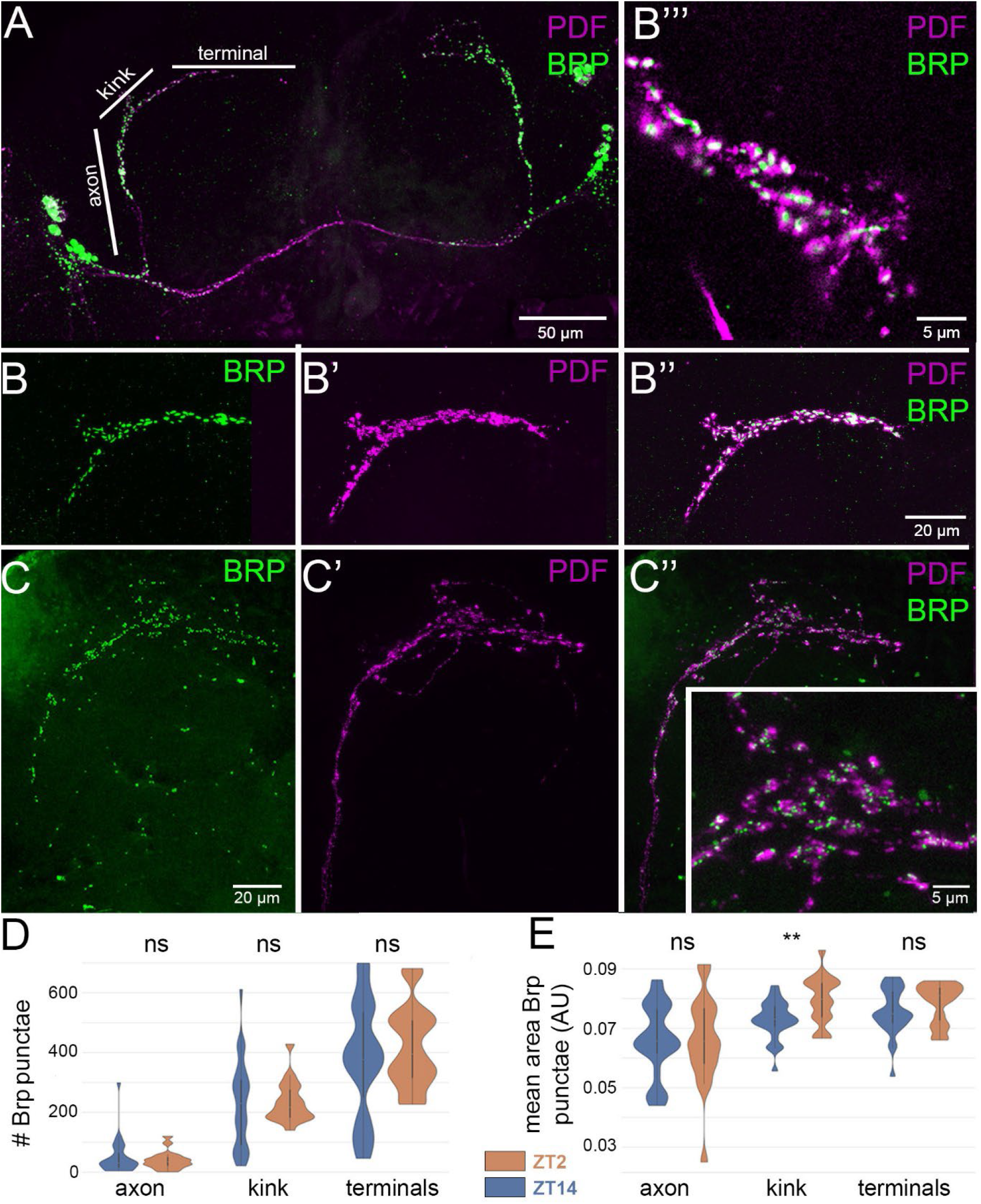
Confocal imaging of BRP and PDF labelling in sLNv. BRP::GFP labelling shown in green, PDF labelling shown in magenta. A) Overview of the LNv morphology and Pdf-Gal4-UAS-driven expression of BRP::GFP. The sLNvs (asterisks) send lateral axons (axon) dorsally to the dorsal protocerebrum. The axon then turns from a lateral to a medial direction in the kink area to eventually arborise in the dorsal terminals in the superior protocerebrum. B-B’’) Close-up of the expression of Pdf-GAL4-UAS-driven expression of BRP::GFP (B) and PDF immunolabeling (B’) in varicosities of the dorsal terminals. B’’) shows the merge of PDF and BRP staining. B’’’) Higher magnification of the dorsal terminals in B’’) shows the accumulation of GFP::BRP in the core of the varicosities, surrounded by PDF staining that labels the sites of peptidergic dense-core vesicles. The cytoplasmic location of GFP::BRP appears to be an artefact due to overexpression by the GAL4-UAS system. C-C’’) Close-up of the expression of endogenously driven expression of BRP^FonYPet^ (dFlex-System) (C) and PDF immunolabeling (C’) in varicosities of the dorsal terminals at the same scale as B-B’’). C’’) shows the merge of PDF and BRP labelling. Insert in C’’) Higher magnification (same scale as in B’’’)) of the dorsal terminals shows small BRP-labelled punctae at the membrane of the varicosities, separated from the more cytoplasmic PDF staining.

We therefore switched to the dFLEx system (Gärtig *et al*., 2019) and expressed fluorescently labelled BRP^FOnYPet^ at endogenous levels in the sLNvs. Using dFLEX, we observed labelling of membrane-associated puncta in a size range around 400 nm in confocal resolution (Fig. 1C) which is compatible with the data from electron microscopic measurements at the NMJ (Kaufmann *et al*., 2002). We obtained similarly sized punctae after Brp^GFP^-labelling using synaptic tagging with recombination (STaR), another method to cell-specifically label presynaptic proteins at endogenous levels (Chen *et al*., 2014) (Suppl. Figure 1). These findings indicate that endogenous labelling by fluorescent BRP constructs reliably reports the size and distribution of active zones in the sLNv. The highest number of BRP^FOnYPet^ punctae was found in the terminals in the superior protocerebrum, followed by the region of the kink in the superior lateral protocerebrum (Fig. 1D). The long proximal axonal stretch from the sLNv somata close to the accessory medulla dorsally to the kink contained much sparser BRP^FOnYPet^ labelling (Fig. 1D). These results are consistent with the reported high density of synaptic input sites in the terminal and kink region, and the sparse occurrence in the proximal axonic region based on the Janelia hemibrain EM dataset (Shafer *et al*., 2022).

### The number of BRP^FOnYPet^-labelled punctae in sLNv terminals remains stable despite circadian remodelling

The results above indicated that the dFLEx system reliably reports the size and location of BRP-labelled active zones in the sLNvs. We next used the dFLEX system to analyse the number of BRP^FOnYPet^ punctae in different areas of PDF-immunostained sLNvs by high magnification confocal microscopy in whole-mount preparations at ZT2 and ZT14. At these times, the sLNv terminals in the dorsal protocerebrum show a high (ZT2) or low (ZT14) degree of branching (Fernández *et al*., 2008; Gorostiza *et al*., 2014). Despite the considerable daily morphological changes of the sLNv terminals, we were unable to find significant differences in the number (Mann-Whitney p= 0.377, Fig 1D)) and mean area (Mann-Whitney p= 0.151, Fig. 1E) of BRP^FOnYPet^ punctae in the terminal region between ZT2 (n= 29 images from 17 brains) and ZT14 (n=18 images from 11 brains) in (Fig. 1D-E) based on high-magnification confocal microscopy. Similarly, the number (Mann-Whitney p= 0.486, Fig 1D)) and mean area (Mann-Whitney p= 0.495, Fig. 1E) of BRP^FOnYPet^ punctae in the proximal axon region, and their number in the kink region (Mann-Whitney p= 0.368, Fig 1D)) did not change significantly between ZT2 and ZT14 in the same preparations (Fig. 1D-E). However, we observed a significantly larger mean area in the kink region at ZT14 (Mann-Whitney, p<0.01, Fig. 1E).

### The density of BRP^FOnYPet^- and PDF-labelled punctae in sLNv terminals changes with circadian remodelling

The unchanged number of BRP^FonYPet^-labelled punctae between ZT2 and ZT14 in the sLNv is not in direct agreement with a previous study (Gorostiza *et al*., 2014). As the size of active zones is typically below 70 nm, we wondered whether these results may be caused by insufficient spatial resolution of confocal light microscopy. We therefore repeated the analysis with expanded brains using a previously established expansion microscopy (ExM) protocol (Gärtig *et al*., 2019). Unfortunately, this protocol resulted in extremely low staining intensities, which would require an optimised custom-made microscopic setup to visualise BRP and PDF staining in the sLNv terminals (Fig. 2A). To circumvent this issue, we performed an additional post-expansion staining which allowed us to image BRP and PDF staining in the sLNv terminals with a standard confocal microscope equipped with sensitive HyD detectors (Fig. 2B-C). The subsequent quantification of the number of BRP^FonYPet^- and PDF-labelled punctae per area in the imaged terminal portions revealed a significant increase in the mean density of BRP^FonYPet^-labelled puncta (Mann-Whitney p<0.001) from ZT2 to ZT14 (Fig. 2D), with the number of punctae more than doubled from ZT2 to ZT14 (from 3.9±0.5 s.e.m. (n=61 images from 7 brains) to 8.4±0.6 s.e.m. (n= 32 from 4 brains). Of note, the overall numbers are about 10x as high as reported earlier (Gorostiza *et al*., 2014), likely due to the increased spatial resolution of ExM combined with marker expression at endogenous levels. Furthermore, the mean density of PDF puncta increased between ZT2 to ZT14 (Fig. 2C-C’, E’), yet to a less pronounced extent (from 14.0±0.7 s.e.m to 15.3±0.9 s.e.m. per µm^2^, Mann-Whitney p<0.05) than for BRP. We further noted that the mean area of the PDF punctae was slightly enlarged by 5% at ZT14 (Mann-Whitney p<0.05, Fig. 2E’). A similar trend was observed for BRP^FonYPet^ punctae (13%, Mann-Whitney p=0.065, Fig. 2D’). Whether this slight increase is of biological relevance or is caused by close contact between separated puncta at the time of the strongest terminal contraction remains undetermined.

**Figure 2:**
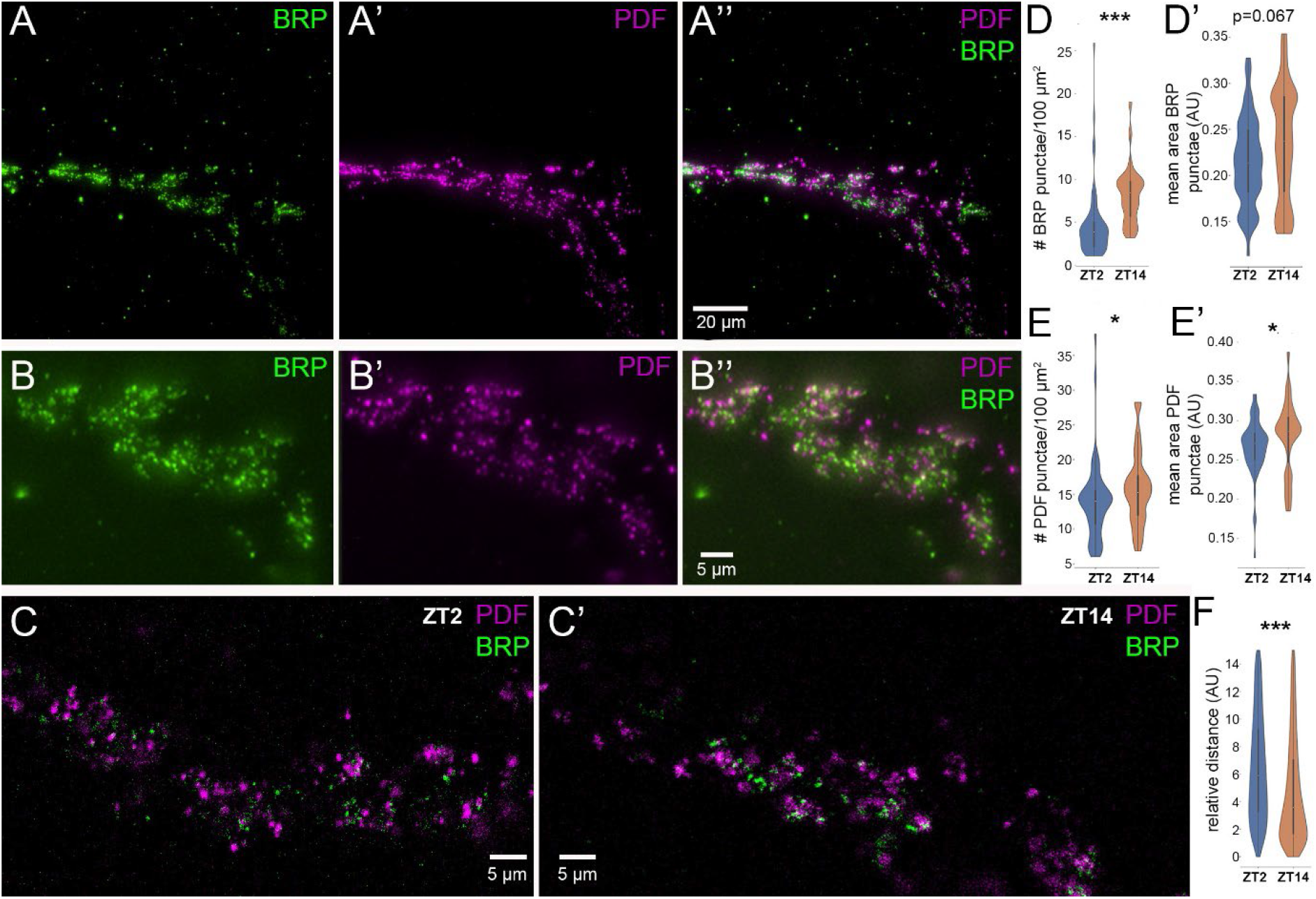
Expansion microscopy of BRP- and PDF-labelled punctae in sLNv terminals. A-A’’) Distribution of BRP^FonYPet^ (A) and PDF-labelled (A’) punctae in the dorsal sLNv terminals, whole stack imaged by a custom-build spinning disc confocal system (Evers lab). B-B’’) Distribution of BRP^FonYPet^ (A) and PDF-labelled (A’) punctae in the dorsal sLNv terminals, whole stack imaged by a confocal laser scanning system. Independent of the confocal system, single punctae can be differentiated after expansion. C-C’) Optical section through a sLNv terminal at ZT2 (C) and ZT 14 (C’). The different punctae appear more concentrated at ZT14. D-D’) Quantification of the number (D) and relative area (D’) of BRP^FonYPet^ punctae per expanded whole stack area in the terminal region suggest a strongly increased active zone density at ZT14, when the terminals are contracted, compared to ZT2 when the terminals show maximum branching. E-E’) Quantification of the number (E) and relative area (E’) of PDF-labelled punctae per expanded whole stack area in the terminal region suggest a slightly increased density at ZT14, when the terminals are contracted, compared to ZT2 when the terminals show maximum branching. F) The relative distance between BRP^FonYPet^ and PDF punctae is strongly decreased because of the increased density. Colour code: BRP^FonYPet^ (green), PDF (magenta). Scales apply to the expanded brain.

In line with the increased density, the mean distance between BRP^FOnYPet^ and PDF punctae was significantly reduced by 40% at ZT14 compared to ZT2 (Mann-Whitney p<0.001, Fig. 2F). The simplest explanation for these findings is a compaction of active zones and PDF-containing vesicles in the retracted terminals at ZT14. Unfortunately, we are unable to give precise distances as the weak fluorescence in the expanded samples did not allow a pre-expansion scan to calculate the degree of expansion anisotropy. Collectively, our data shows that the number of BRP-organised active zones in the sLNv terminals stays fairly stable despite considerable morphological changes in the dorsal sLNv terminals between ZT2 and ZT14.

### Lack of spatial correlation between spontaneous peptidergic vesicle release and active zones

To independently test for a lack of spatial correlation between sites of dense-core vesicle release and active zones, we relied on the available dataset of an entire female adult fly brain reconstructed from serial section transmission electron microscope (ssTEM) slices (FAFB, https://fafb.catmaid.virtualflybrain.org/, https://temca2data.org) (Zheng *et al*., 2018). We first identified in total seven neurons (4 in the left, 3 in the right hemisphere) that have the unique sLNv morphology and contain dense-core vesicles and reconstructed them using CATMAID (Suppl Fig. 2). We note that this morphological match does not necessarily prove the identity of these putative sLNvs in the FAFB brain. Yet, to the best of our knowledge, there is no other neuron type with similar anatomy. Further, the s-LNvs from the hemibrain (Shafer *et al*., 2022) transformed to the FAFB dataset fit the location and arborisation pattern of the identified neurons. In addition, the cell numbers match well (each hemisphere contains four sLNvs (Helfrich-Förster, 1997)). We therefore concluded that these neurons are sLNvs, and went through each serial section to look for spontaneous release profiles (“Ω profiles”) of dense-core vesicles along the proximal axon, the kink and the terminal region that contains the vast majority of synaptic outputs (Shafer *et al*., 2022). In total, we identified 58 Ω profiles and 89 docked dense-core vesicles which were quite evenly distributed along the proximal axon, kink and terminal region (Fig. 3A), unlike the previously described presynaptic sites of the sLNvs in the hemibrain that are clustered in the terminal and a few zones along the proximal axon (Shafer *et al*., 2022). None of the Ω profiles and docked dense-core vesicles was close to active zones (Fig. 3B-G). While dense-core vesicles close to synaptic vesicle pools occurred, we never observed fusion or docking of the dense-core vesicles in the immediate vicinity of these pools close to active zones. Collectively, the analysis of the FAFB ssTEM dataset supports the independence of peptidergic dense-core vesicle release from active zones and presynaptic regions of the sLNvs.

**Figure 3:**
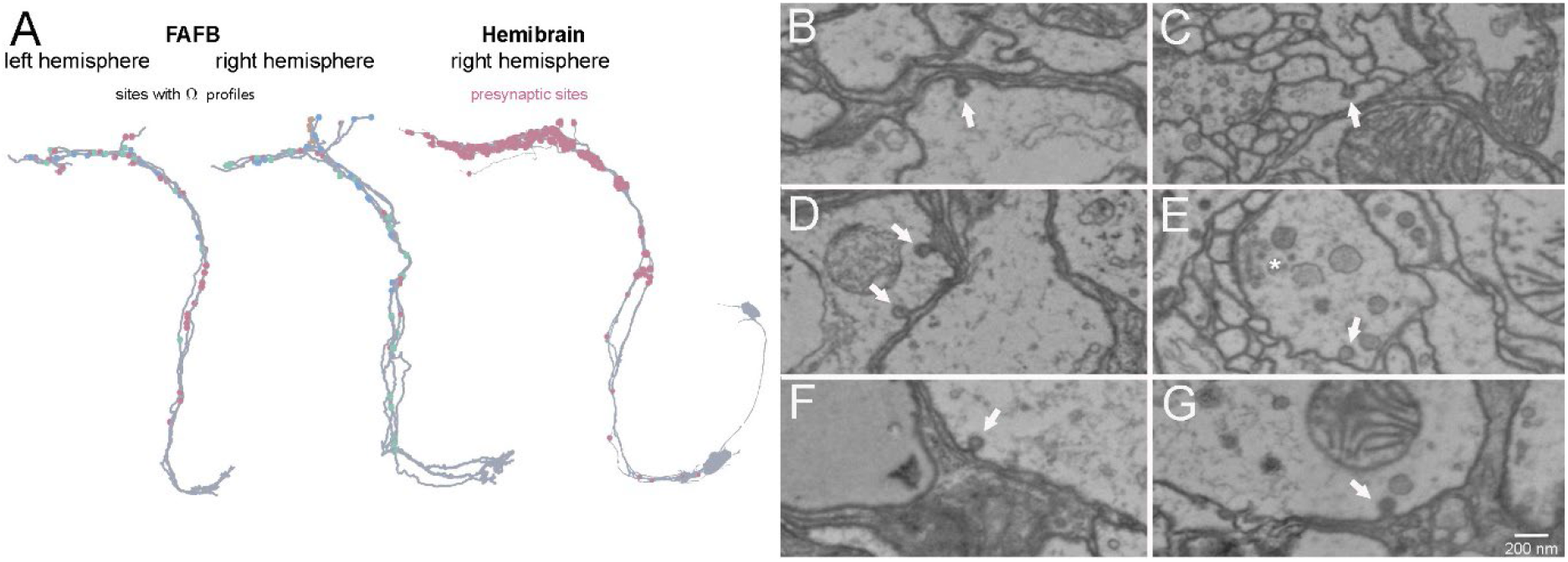
Examples of dense-core vesicle fusion (Ω profiles) in the sLNv in the FAFB brain. A) Mapping of identified Ω profiles to the FAFB sLNv skeletons (left), and comparison to the distribution of identified presynaptic sites in the sLNvs from the hemibrain (hembibrain data from (Shafer et al., 2022)). The Ω profiles are more evenly distributed in the ascending primary neurite than the presynaptic sites which are clustered to a few spots along the primary neurites. In the terminals, presynaptic sites but not Ω profiles are densely clustered. Ω profiles are colour-coded according to single sLNvs (see Suppl.Fig.2). B-G) Single TEM sections with fusing dense-core vesicles (arrows) in different anatomically identified sLNv. All Ω profiles observed were distant to active zones (also in the z-axis, not shown). E) shows an example where an active zone with synaptic vesicle pool (asterisks) is seen in the same section. Scale bar in G) applies to all sections.

### The sLNvs express transcripts for BRP and other active zone proteins

Next, we used a bioinformatics approach to determine which genes coding for a targeted set of synaptic proteins are expressed in the sLNvs, relying on available single-cell RNAseq data of clock neurons from the Rosbash lab (Ma *et al*., 2021). sLNvs were identified and differentiated from the large LNvs (lLNvs) based on the co-expression of PDF, sNPF and AstC-receptor 2 (Ast-C R2), while the lLNvs express PDF but neither sNPF nor Ast-C R2 (Johard *et al*., 2009; Ma *et al*., 2021) (Fig. 4A-B). We found all targeted active zone and SNARE protein genes to be expressed in both the small and large LNvs (Fig. 4C), along with the gene coding for Calcium-dependent secretion activator (*Cadps*), a protein required for synaptic- and dense-core vesicle exocytosis (Renden *et al*., 2001). We further found expression of *amon*, a gene that encodes the prohormone convertase dPC2 required for posttranslational neuropeptide processing (Wegener *et al*., 2011). Transcript expression for the targeted synaptic proteins seems to occur independent of light input, as the expression levels were similar between LD and DD conditions (Fig. 4D). While diel or circadian changes in expression levels are visible for all analysed synaptic proteins, the various transcripts are not synchronised with each other and do not cycle in phase with Pdf transcripts (Fig. 4E-F). *Pdf* expression is highest from ZT22-ZT06/CT22-CT02. This is in line with a significantly higher number of PDF puncta at ZT2 compared to ZT14, and PDF release during the early light phase (Park *et al*., 2000; Klose *et al*., 2021) when fast Ca^2+^ activity peaks in the “morning neurons” (Liang *et al*., 2022). Yet, it is out of phase with *brp* expression which is highest during ZT14-18/CT14-22 and (Fig. 4E-F). This temporal mismatch provides additional support for a BRP-independent exocytosis of PDF.

**Figure 4:**
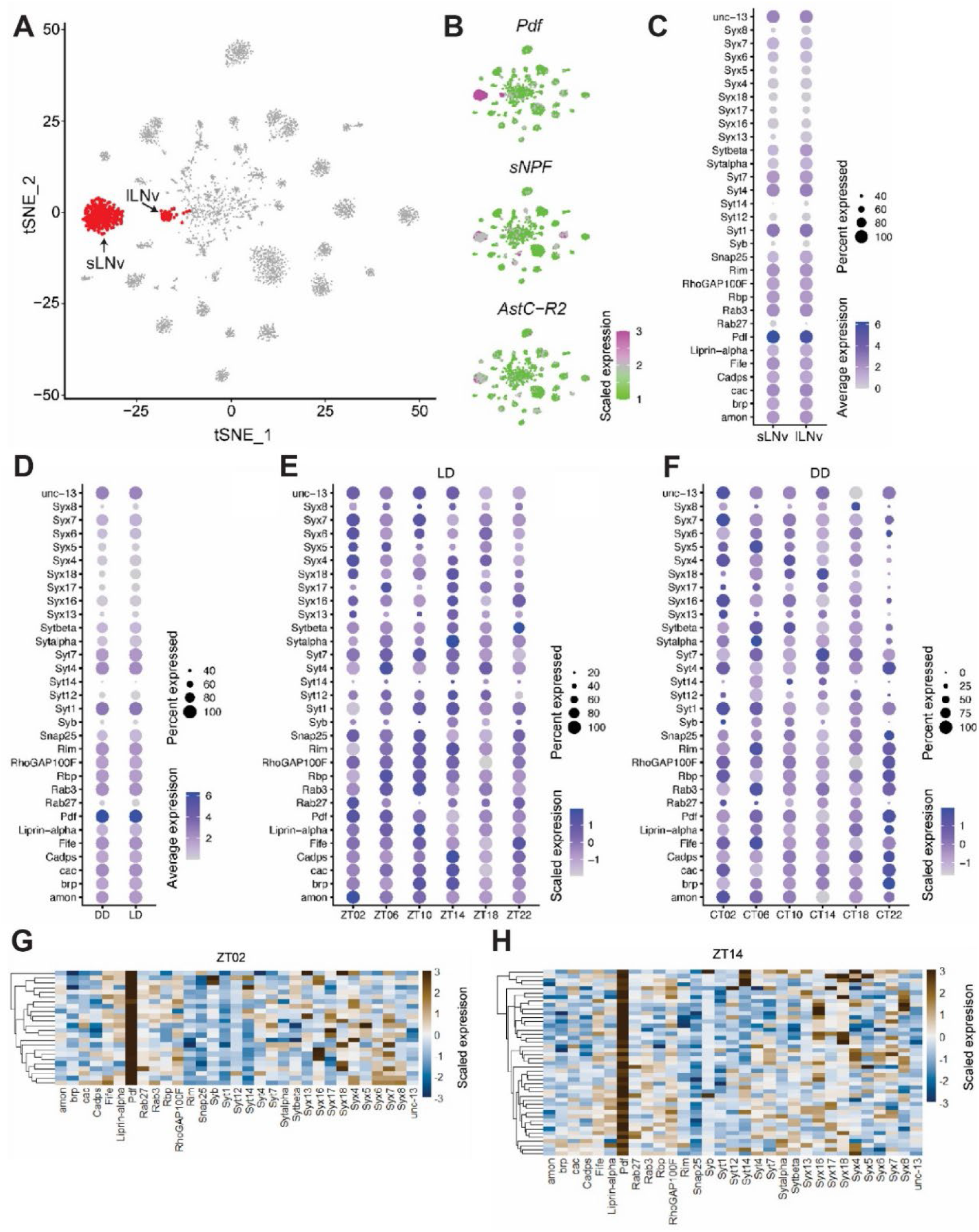
Single-cell transcriptome analysis of clock neurons. A) A tSNE plot of clock neuron (Clk856-GAL4) single-cell transcriptomes showing large lateral ventral neuron (lLNv) and small lateral ventral neuron (sLNv) cell clusters. B) tSNE plots showing expression of pigment dispersing factor (Pdf), small neuropeptide F (sNPF) and allatostatin-C receptor 2 (AstC-R2). Pdf is highly expressed in both lLNVs and sLNVs but sNPF and AstC-R2 are expressed in sLNvs but not lLNvs. C) Dot plot showing expression of genes in lLNvs and sLNvs. D-F) Expression of genes under light-dark conditions (LD – 12h light:12h dark) and under constant darkness (DD). D) Dot plot comparing expression of genes under DD and LD. E) Dot plot comparing expression of genes at six different timepoint under LD conditions. F) Dot plot comparing expression of genes at six different timepoint under DD conditions. G-H) Comparison of gene expression at specific timepoints reveals sLNv heterogeneity. Heatmaps showing expression of genes at (G) Zeitgeber 2 (ZT2) and (H) ZT14. Each row represents a single cell. Pdf is highly expressed in all cells, but expression of other genes is variable between different cells and across different time points.

### CADPS and SNARE proteins, but not BRP seem to be required for PDF signalling

The data above provided evidence that PDF dense-core vesicle release is independent of BRP-organised active zones. To test whether BRP and associated active zone proteins are functionally required for PDF release, we performed a *Pdf-*Gal4-driven RNAi screen against a subset of the targeted synaptic and exocytosis-related genes found to be expressed in the sLNvs. Rhythmic locomotor activity was chosen as behavioural read-out, taking advantage of the high level of arrhythmic locomotor activity in DD in flies with a null mutation in the *Pdf* gene (*Pdf^01^,* (Renn *et al*., 1999)). *Pdf^01^* mutants that remain rhythmic in DD show rather weak locomotor rhythmicity with significantly shorter periods (Renn *et al*., 1999). *Pdf^01^* mutants are further unable to delay their evening activity during long photoperiods (Yoshii *et al*., 2009) which prompted us to use long day conditions (LD16:8). *Pdf^01^* mutant flies and RNAi lines against *Pdf* and *amon* were included as controls. The results are summarised in Table 1. As expected, *Pdf^01^* and *Pdf-*RNAi flies became arrhythmic in DD (Fig. 5A-B). Surprisingly, down-regulation of *amon* did not result in a significantly altered rhythmicity, although it has been shown to be highly efficient in other peptidergic cells (Rhea *et al*., 2010). This finding suggests that substantial knock-down of PDF signalling is required to significantly impair rhythmicity. *Cadps*-RNAi^110055^ resulted in a strong arrhythmic phenotype, similar to complete loss or knock-down of *Pdf*. RNAi against the transcripts of the SNARE protein-coding genes *Syntaxin* (*Syx*) 1, *Syx4*, *Syx18,* and *neuronal Synaptobrevin* (*nSyb*), as well as *Rbp* and *unc13A* resulted in <50% rhythmic flies and a significantly (p<0.01) reduced rhythmic strength compared to *Pdf*-Gal4 controls (Fig. 5A-B). *Rab3*-RNAi^100787^ flies showed significantly reduced rhythmic strength, while 58% remained rhythmic. In contrast, none of the RNAi constructs against *brp*, S*yd-1 (RhoGAP100F)*, *liprin-*α*, RIM*, *fife*, *unc13B, cac,* as well as the SNARE gene *SNAP25* affected rhythmicity. Collectively, our results suggest that vesicle-associated membrane proteins (CADPS, nSYB) and their direct cell membrane-associated partner SYX are essential parts of the machinery for PDF release. Furthermore, active zone scaffold proteins like BRP, SYD1, and LIPRINα seem not to be essential. A main function of BRP at the NMJ is to localize voltage-gated channels including the α1 subunit isoform CACOPHONY (CAC) at the active zone (Fouquet *et al*., 2009); thus the lack of effect of *cac^RNAi^* on locomotor rhythmicity provides further evidence against an important role of assembled active zones for PDF release.

**Figure 5:**
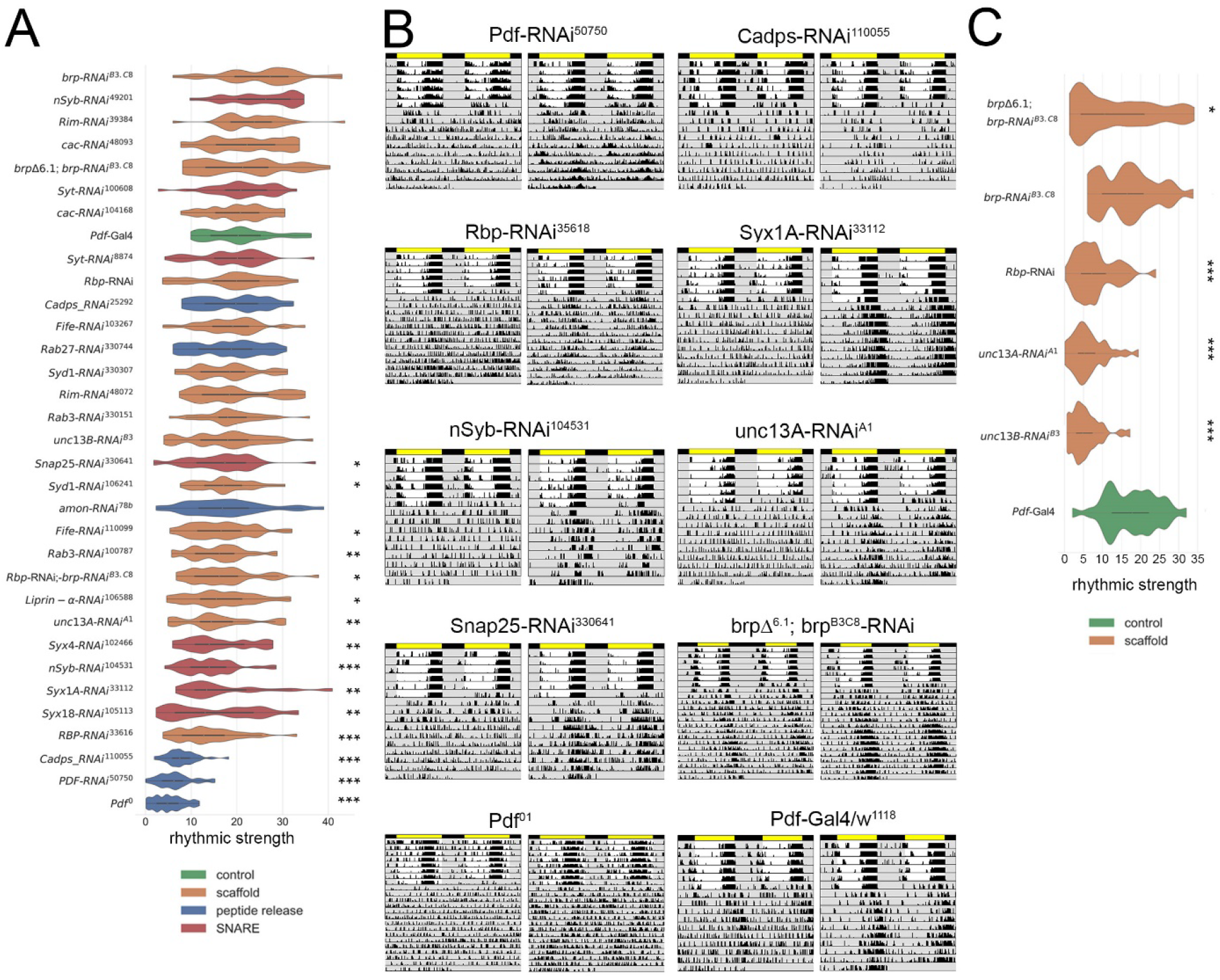
Locomotor activity and rhythmicity after RNAi-mediated knock-down of synapse- and vesicle-release associated genes. A) Autocorrelation rhythmicity strength of circadian locomotor activity in DD after RNAi-mediated knockdown driven by *Pdf*-Gal4. The strongest reduction in rhythmicity was found for Pdf^0^ mutants, and after knock-down of *Pdf* and *Cadps*. A significant reduction in rhythmicity was also achieved by knock-down of several SNARE and active zone scaffold genes, but not by various *brp* RNAi lines. B) Example actograms of individual flies after gene knock-down with the indicated RNAi lines, and in *Pdf^01^*mutants and *w^1118^* controls. The left actogram shows the least rhythmic, the right actogram the most rhythmic fly in the measured series. C) Autocorrelation rhythmicity strength of circadian locomotor activity in constant light (LL, 1 Lux). A strong reduction in rhythmicity after knock-down of various active zone-associated genes including *brp* is visible under these conditions uncovering glycine transmission.

**Table 1:**
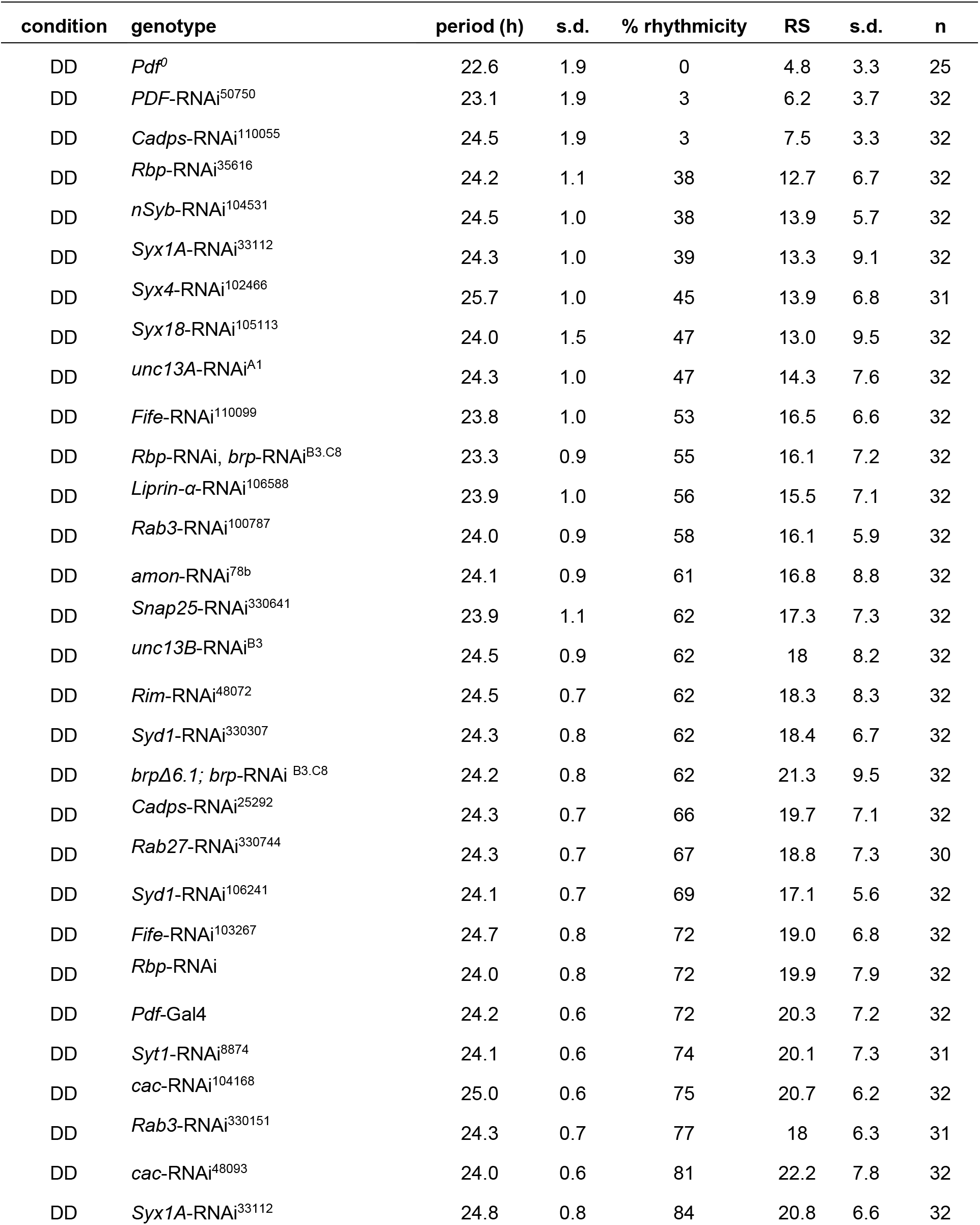
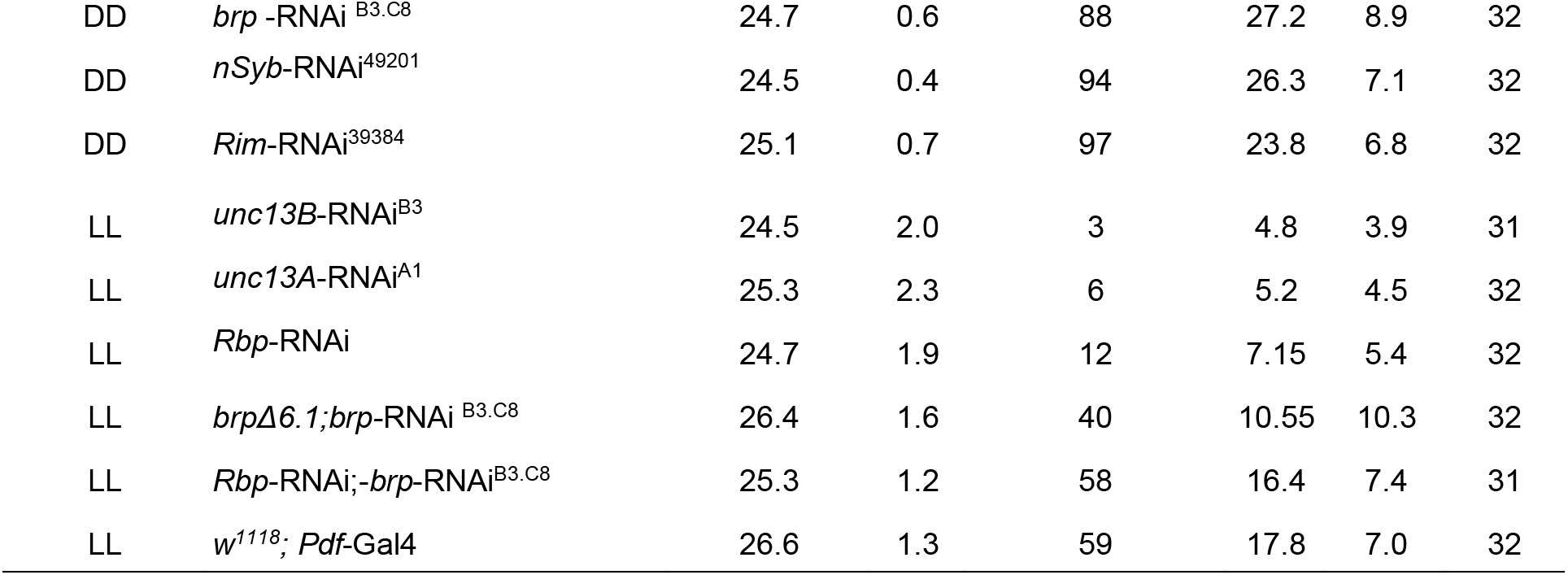
Locomotor rhythmicity of flies with RNAi-mediated knock-down of synapse-associated proteins and controls. Period, rhythmicity, and rhythmic strength (RS) derived from autocorrelation analysis. For genotypes see Suppl. Table 1.

We next downregulated transcripts of core active zone scaffold proteins by RNAi in the sLNv and monitored locomotor rhythmicity under constant dim light (LL, 1 lux). Under these conditions, the majority of flies impaired in signalling becomes arrhythmic (Frenkel *et al*., 2017). Glycine is a classic transmitter co-localised with PDF in the sLNv known to be released via small synaptic vesicles at active zones. We therefore reasoned that a good behavioural readout to test whether RNAi inefficiency against active zone scaffold proteins might underlie the absent effect of RNAi against *Brp* and other active zone genes under DD. In our hands, 59% of *Pdf*-Gal4 controls remained rhythmic in LL, while knock-down of *unc*13A, *unc*13B, *Rbp*, and *brpΔ6.1; brp-*RNAi ^B3.C8^ led to a significantly lowered rhythmicity (Fig. 5C), likely by impairing the release of glycine via small synaptic vesicles. This finding suggests that the lack of effect of RNAi *Brp* and other active zone scaffold proteins on PDF-dependent rhythmicity is not due to inefficient transcript down-regulation. However, *Rbp*-RNAi; b*rp*-RNAi^B3.C8^ had no effect, suggesting that a strong knock-down of *Rbp* and *brp* is required to impair glycine release.

## Discussion

An impressive body of knowledge exists on the terminal synaptic plasticity and circadian release of PDF from central sLNv clock neurons, as outlined in the introduction. However, the extent to which circadian peptide release and terminal and synaptic plasticity in sLNvs are mechanistically linked to each other is ill-defined. While some researchers speculated whether rhythmic PDF levels may be a secondary consequence of circadian remodeling of the sLNv terminals (King & Sehgal, 2018) or used PDF as a marker of presynaptic boutons (Bushey *et al*., 2011), ultrastructural data suggested that PDF-containing DCV are non-synaptically released (Yasuyama & Meinertzhagen, 2010).

Here we provide combined microscopic and neurogenetic evidence for an independence of PDF release from BRP-labelled active zones in the sLNvs. First, we were unable to find a spatial and numerical correlation between PDF-labelled dense-core vesicles and BRP-labelled active zones using super-resolution microscopy. In addition, we found that the temporal pattern of PDF and BRP mRNA expression is not in phase with each other. Further, our RNAi screen points to CADPS and a set of SNARE proteins, but not BRP or other active zone scaffold proteins as being essential for peptide release from the sLNvs. On the ultrastructural level, we found that the spontaneous sites of DCV release across the sLNv are not restricted to certain neuronal compartments and are consistently well separated from active zones, in line with earlier EM studies on sections of the sLNv terminals (Yasuyama & Meinertzhagen, 2010) or the terminal arborisations of the large PDF-expressing LNv in the visual system (Miśkiewicz *et al*., 2008). Collectively, these results show that PDF release is independent of BRP-organised active zones.

An unexpected finding of our study was the obvious constancy of the number of BRP punctae between the times of day with maximum (ZT2) and minimum (ZT14) branching of sLNv terminals. This should lead to an increased density of BRP punctae from ZT2 to ZT14 which was what we observed. Our data suggest that BRP is not cyclically degraded and produced, but rather changes its location; however, this requires further investigation. Interestingly, a similar compaction of numerically unchanged BRP on a smaller mesoscale has been recently described for the *Drosophila* neuromuscular junction during presynaptic homeostatic potentiation (Mrestani *et al*., 2021; Dannhäuser *et al*., 2022; Ghelani *et al*., 2023). In the lamina, however, the number and size of presynaptic profiles as well as the level of BRP are changing during BRP-dependent circadian plasticity (Górska-Andrzejak *et al*., 2013; Górska-Andrzejak *et al*., 2015; Woźnicka *et al*., 2015). While our results do not add mechanistical insight here, it seems possible that the mechanism of circadian plasticity and the dependence on BRP differ between the sLNvs and lamina.

To functionally support the notion of a BRP-independent PDF release, we systematically downregulated transcripts of *brp* and related genes in the PDF neurons and tested whether this mimics a loss of PDF signalling. Though in general powerful, RNAi-mediated downregulation is bound to potential caveats. The RNAi downregulation may not be strong enough or have off-target effects. We therefore used at least two different RNAi constructs where possible. Second, and perhaps more severe, there is a considerable degree of mutual redundancy among several active zone proteins (Ghelani & Sigrist, 2018; Held & Kaeser, 2018). To account for this, we used combinations of different RNAi constructs against BRP, performed downregulation in a heterozygous *brp* deficient background, and also performed simultaneous knockdown of *brp*- and *Rbp* that was previously developed and successfully used by the Sigrist group (Wagh *et al*., 2006; Fouquet *et al*., 2009; Petzoldt *et al*., 2020). None of these combinations affected PDF-dependent locomotor activity in DD, but at least *brp*Δ6.1; *brp*-RNAi^B3.C8^ impaired glycine-dependent locomotor activity in LL. We interpret these results as functional support for a BRP-independence of PDF release from the sLNvs. A role for ELKS in peptide release has been described for insulin secretion in pancreatic β-cells (Ohara-Imaizumi *et al*., 2019). Yet, peptidergic ecdysis-triggering hormone (ETH) release from the endocrine epitracheal cells also appeared to be BRP-independent, though BRP is expressed by those cells (Bossen *et al*., 2023).

The results obtained after downregulation of other active zone or vesicle release-related genes are difficult to interpret as in some cases only one of the two employed RNAi constructs gave effects. Moreover, we did not test different combinations of RNAi constructs. CADPS stands out from this, as downregulation of its transcripts produced a strong PDF-related phenotype. *Cadps* RNAi was also very effective in downregulating insulin (DILP7) (Imambocus *et al*., 2022) and ETH release (Bossen *et al*., 2023) in *Drosophila*. Thus, our results support the requirement of CADPS for dense-core vesicle release as has been previously reported for the fly and mammals (Renden *et al*., 2001; Farina *et al*., 2015). Our results also support the role of SNARE proteins in dense-core vesicle release (Chen & Scheller, 2001; Hoogstraaten *et al*., 2020) and provide ambiguous evidence for the requirement of RIM which has been shown to be essential for dense-core vesicle release in mammalian neurons (Persoon *et al*., 2019).

Taken together, our results based on anatomy and a functional RNAi screen infer that PDF release from the sLNvs is independent of BRP but requires CADPS and the canonical secretory vesicle fusion machinery. As BRP is a key organiser of active zones in *Drosophila* (Wagh *et al*., 2006; Ghelani & Sigrist, 2018), we further assume that PDF release is likely independent of BRP-organised active zones and, in principle, can occur along most axon-derived processes in a non-localised fashion. This conclusion is supported by our finding of a relatively even distribution of spontaneous peptide release events along the entire projections of the sLNvs in the FAFB brain. Moreover, a previous study showed PDF release from the soma of sLNvs (Klose *et al*., 2021), a compartment lacking active zones. A BRP/active zone-independent and non-localised release of PDF would also explain the mild circadian phenotype of flies with genetically abrogated sLNv terminals (Fernandez *et al*., 2020). From a generic perspective, active zone-independent release of peptidergic dense-core vesicles seems to be the rule rather than the exception (Ludwig & Leng, 2006; Salio *et al*., 2006; Nässel, 2009), and has been ultrastructurally described already more than 40 years ago across bilaterian taxa, including molluscs (Buma & Roubos, 1986), crayfish (Schürmann *et al*., 1991) and rats (Pow & Morris, 1988, 1989). One possibility how PDF-containing dense-core vesicles could achieve non-localised release is via recruitment of SNARE proteins such as SYX (Knowles *et al*., 2010; Gandasi & Barg, 2014). This needs further testing but we note that transcriptional down-regulation of SYX and other SNARE proteins significantly impaired PDF signalling from the LNvs in our study. Our results further suggest that BRP is unlikely to be broken down and reassembled and synthesized again during circadian plasticity. We propose that a seemingly simpler relocation may provide a more efficient and cost-effective way of synaptic plasticity.

## Author contributions

Conceptualisation: BH, CWeg, with input from JFE and MZ. Labelling and imaging: BH, JFE, CWer. Behaviour: BH, DR. CATMAID and image analysis: BH, NR. scRNAseq analysis: MZ. Supervision: CWeg, JFE. Funding acquisition: CWeg. Wrote the manuscript draft: BH, CWeg, with input from MZ and NR. All authors read and commented on the final draft of the manuscript.

## Acknowledgment

We thank Markus Kiunke for setting up the server and preparing the dataset for analysis, Stephan Sigrist and David Owald for helpful discussions, and Michael Bender, Stephan Sigrist, and Paul Taghert for the kind gift of flies. We especially thank Marta Costa and the FAFB community for creating the dataset and making it open source. Funding was provided by a programme of the Faculty of Biology, JMU (to CW) and the Deutsche Forschungsgemeinschaft (DFG, German Research Foundation): 251610680, INST 93/809-1 FUGG, for the Leica TCS SP8 microscope).

## Data availability statement

Upon publication, the data will be made publicly available at WUEDATA, the JMU research data repository.

