## Supplemental Table and Figures for "PDF neuropeptide signals independently of Bruchpilot-labelled active zones in daily remodelled terminals of *Drosophila* clock neurons"

### Supplementary Material

**Supplementary Table 1:** Fly lines used for the RNAi screen.

| Genotype | Source | library/# | CG | chromosome | reference |
|---|---|---|---|---|---|
| **UAS-RNAi lines** | | | | | |
| UAS-*amon*-RNAi$^{78b}$ | M. Bender | | 6438 | 3 | [1] |
| UAS-*brp* -RNAi $^{B3.C8}$ | S. Sigrist | | 42344 | 3 | [2] |
| UAS-*brp*-RNAi $^{B3}$ | S. Sigrist | | 42344 | 3 | [3] |
| UAS-*brp*-RNAi$^{C8}$ | S. Sigrist | | 42344 | 3 | [3] |
| *brpΔ6.1;brp*-RNAi $^{B3.C8}$ | S. Sigrist | | 42344 | 2;3 | [2] |
| UAS-*cac*-RNAi$^{104168}$ | VDRC | KK | 43368 | 2 | |
| UAS-*cac*-RNAi$^{48093}$ | VDRC | GD | 43368 | 1 | |
| UAS-*Cadps*-RNAi$^{110055}$ | VDRC | KK | 33653 | 2 | |
| UAS-*Cadps*-RNAi$^{25292}$ | VDRC | GD | 33653 | 3 | |
| UAS-*Fife*-RNAi$^{103267}$ | VDRC | KK | 43955 | 2 | |
| UAS-*Fife*-RNAi$^{110099}$ | VDRC | KK | 43955 | 2 | |
| UAS-*GlyT*-RNAi$^{330127}$ | VDRC | shRNA | 5549 | 2 | |
| UAS-GlyT-RNAi$^{8222}$ | VDRC | GD | 5549 | 2 | |
| UAS-*Liprin-α*-RNAi$^{106588}$ | VDRC | KK | 11199 | 2 | |
| UAS-*Liprin-α*-RNAi$^{51707}$ | VDRC | GD | 11199 | 2 | |
| UAS-*nSyb*-RNAi$^{104531}$ | VDRC | KK | 17248 | 2 | |
| UAS-*nSyb*-RNAi$^{49201}$ | VDRC | GD | 17248 | 3 | |
| UAS-*Pdf*-RNAi$^{50750}$ | VDRC | GD | 6496 | 3 | |
| UAS-*Rab27*-RNAi$^{330744}$ | VDRC | shRNA | 14791 | 2 | |
| UAS-*Rab3*-RNAi$^{100787}$ | VDRC | KK | 7576 | 2 | |
| UAS-*Rab3*-RNAi$^{330151}$ | VDRC | shRNA | 7576 | 2 | |
| UAS-*Rbp*-RNAi | S. Sigrist | | 43073 | 2 | [4] |
| UAS-*Rbp*-RNAi$^{35616}$ | VDRC | GD | 43073 | 3 | |
| UAS-*Rbp*-RNAi; UAS-*Brp*-RNAi$^{B3.C8}$ | S. Sigrist | | | 3 | |
| UAS-*Rim*-RNAi$^{39384}$ | VDRC | GD | 33547 | 2 | |

| | | | | | |
|---|---|---|---|---|---|
| UAS-*Rim*-RNAi$^{48072}$ | VDRC | GD | 33547 | 3 | |
| UAS-*Snap25*-RNAi$^{330641}$ | VDRC | shRNA | 40452 | 2 | |
| UAS-*c* | VDRC | KK | 1976 | 2 | |
| UAS-*Syd1*-RNAi$^{330307}$ | VDRC | shRNA | 1976 | 2 | |
| UAS-*Syt1*-RNAi$^{100608}$ | VDRC | KK | 3139 | 2 | |
| UAS-*Syt1*-RNAi$^{8874}$ | VDRC | GD | 3139 | 3 | |
| UAS-*Syt1*-RNAi$^{8874}$ | VDRC | GD | 3139 | 3 | |
| UAS-*Syx1A*-RNAi$^{33112}$ | VDRC | GD | 31136 | 3 | |
| UAS-*Syx4*-RNAi$^{102466}$ | VDRC | KK | 2715 | 2 | |
| UAS-*Syx18*-RNAi$^{105113}$ | VDRC | KK | 13626 | 2 | |
| UAS-*unc13A*-RNAi$^{A1}$ | S. Sigrist | | 2999 | 3 | [5] |
| UAS-*unc13B*-RNAi$^{B3}$ | S. Sigrist | | 2999 | 3 | [5] |
| **Gal4 driver line** | | | | | |
| *Pdf*-Gal4 | P. Taghert | | 6496 | 2 | [6] |
| combined with: | | | | | |
| UAS-*dcr* | VDRC | 60007 | | 1 | |
| **Controls** | | | | | |
| *Pdf*$^{01}$ | P. Taghert | | 6496 | | [6] |
| *w*$^{1118}$ | own coll. | | | | |

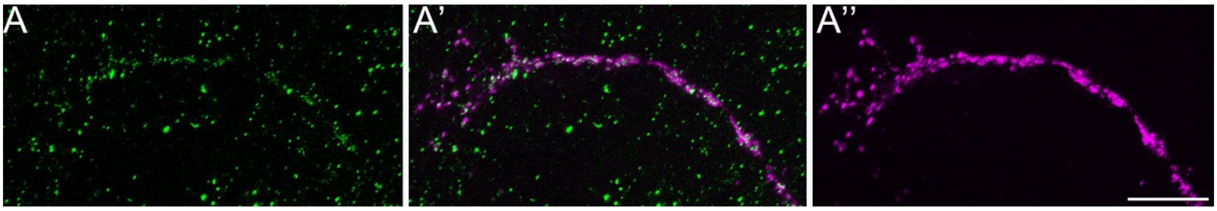


***Supplementary Figure 1*: *Confocal imaging of BRP and PDF labelling in sLNv*.** *A) BRP$^{GFP}$-labelling by synaptic tagging with recombination (STaR) in Pdf-Gal4/UAS-FLP;+/brp-FRT-stop-FRT-GFP flies. The labelled BRP$^{GFP}$ puncta are much smaller than those obtained after Pdf>brp$^{GFP}$ expression and are similar in size to those observed by dFLEX (see text). The numerous green dots of various size outside the sLNv terminal represent unspecific background labelling. A') Overlay of BRP$^{GFP}$-labelling in (A) and PDF immunolabelling (A''). A'') PDF immunolabeling that marks an sLNv terminal. Scale bar = 20 µm.*

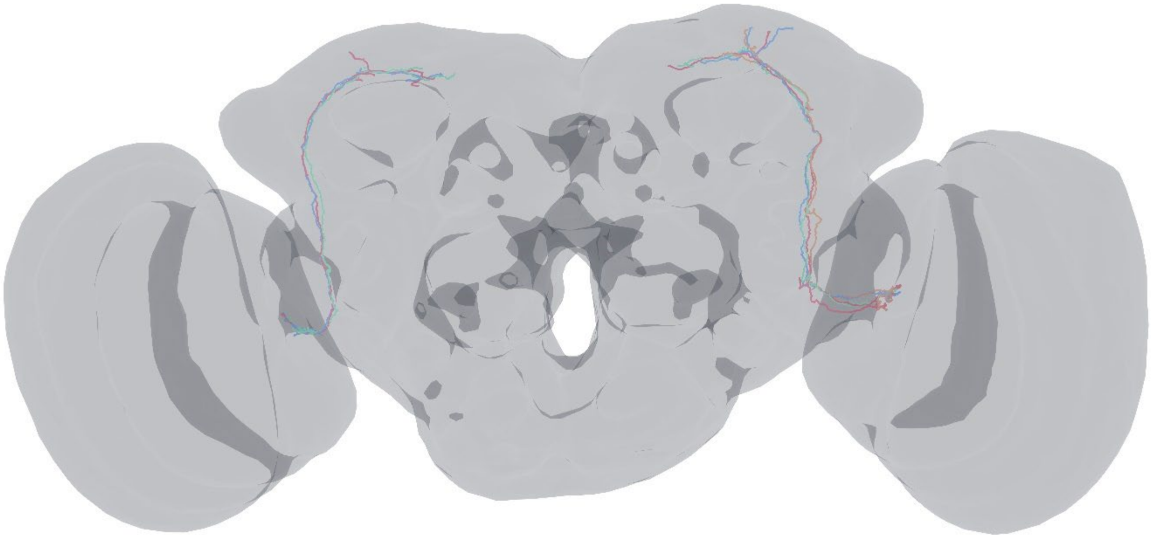

***Supplementary Figure 2:*** *Skeleton reconstructions of four neurons in the left and three neurons in the right FAFB brain hemisphere. The neurons show all hallmarks of the PDF-expressing sLNv including the presence of dense-core vesicles, and somata close to the accessory medulla. To the best of our knowledge, no other peptidergic neurons have a similar anatomy. Therefore, we conclude that these seven neurons are identical to PDF-expressing sLNv. The individual neurons use the same colour code as Fig. 3A.*
